# Impairments in contractility and cytoskeletal organisation cause nuclear defects in nemaline myopathy

**DOI:** 10.1101/518522

**Authors:** Jacob A Ross, Yotam Levy, Michela Ripolone, Justin S Kolb, Mark Turmaine, Mark Holt, Maurizio Moggio, Chiara Fiorillo, Johan Lindqvist, Nicolas Figeac, Peter S Zammit, Heinz Jungbluth, John Vissing, Nanna Witting, Henk Granzier, Edmar Zanoteli, Edna C Hardeman, Wallgren-Pettersson Carina, Julien Ochala

## Abstract

Nemaline myopathy (NM) is a genetically heterogeneous skeletal muscle disorder caused by mutations predominately affecting contractile filaments, in particular thin filament structure and/or regulation. The underlying cellular pathophysiology of this disease remains largely unclear. Here, we report novel pathological defects in skeletal muscle fibres of mice and patients with NM, including disrupted nuclear envelope, altered chromatin arrangement, and disorganisation of the cortical cytoskeleton. We demonstrate that such nuclear defects are caused by impairment of muscle fibre contractility, and that cytoskeletal organisation determines nuclear morphology. Our results overlap with findings in diseases caused by mutations in nuclear envelope or cytoskeletal proteins. Given the important role of nuclear shape and envelope in regulating gene expression, and the cytoskeleton in maintaining muscle fibre integrity, our findings are likely to underlie some of the hallmarks of NM, which include broad transcriptional alterations, arrested muscle fibre growth, contractile filament disarray and altered mechanical properties.

## Introduction

Nemaline myopathy (NM) is a disease of skeletal muscle caused by mutations in genes that are generally involved in muscle contraction, in particular those related to the structure and/or regulation of the thin filament. Mutations in *ACTA1* (skeletal muscle actin) or *NEB* (nebulin) together make up the majority of cases, whilst other causative and as yet unidentified genes are implicated in the remainder (to date, *TPM3*, *TPM2*, *TNNT1*, *CFL2*, *KBTBD13*, *KLHL40*, *KLHL41*, *LMOD3*, *MYPN* or *MYO18B*)^1^. These mutations result in weakness at the contractile level, while other cellular pathological hallmarks include dense accumulations of proteins known as nemaline rods, arrested muscle fibre growth, impaired fibre type differentiation, and disarray of contractile filaments^1^. However, the underlying mechanisms behind these features remain uncertain, even though the mutations affecting thin filament structure and function are likely to be involved^2–9^. In the present study, we aimed to acquire a clearer understanding of muscle fibre dysfunction in NM by specifically studying nuclei and the related cortical cytoskeleton.

Skeletal muscle fibres are large syncytial cells containing many nuclei (termed myonuclei). Sufficient numbers and regular spacing of myonuclei throughout the muscle fibre are a prerequisite for its function, allowing the efficient delivery of gene products to all parts of the cell, with minimal transport distances. Therefore it is thought that each nucleus is responsible for maintaining a certain volume of the muscle fibre, termed the myonuclear domain^10,11^. The nuclei of skeletal muscle fibres are linked with various cytoskeletal components including non-sarcomeric/cytoplasmic actins, microtubules and intermediate filaments such as desmin. All three of these cytoskeletal networks have been implicated in the spacing and positioning of nuclei in models of skeletal muscle development: microtubules in the initial translocation/spacing of nuclei along the fibre^12–15^, and actin and desmin in their movement to the fibre periphery^16^.

At the organelle level, nuclear function and transcription are regulated by a host of external factors; the nuclear envelope acts as a signalling hub that is capable of transducing a range of chemical and mechanical signals to regulate gene expression^17–19^. The cytoskeleton is known to regulate nuclear shape and morphology via interactions with the nuclear envelope^20,21^, a process which can itself impact on gene transcription, with different morphologies being linked to cell type, function, differentiation and disease states^20–22^. Given that the force-generating properties of NM muscle fibres are severely limited, we hypothesised that cytoskeletal components as well as nuclear function, positioning and integrity might be affected in this disease, and possibly contribute to pathology.

Using single muscle fibres from mouse models and NM patients with mutations in *ACTA1* or *NEB*, we found that myonuclei display a range of defects, including irregular spacing, morphological and nuclear envelope abnormalities and altered chromatin distribution. We also observed severe disruption within the microtubule, desmin and cytoplasmic (β- and γ-) actin networks, as well as alterations in their anchorage at the nuclear surface. We next sought to define the underlying mechanisms behind these defects, and found that nuclear spacing and morphology are regulated by contractile force production. We further demonstrated the role of a properly organised microtubule network in regulating nuclear shape. Our findings suggest that these alterations are likely to contribute to some of the features observed in NM, which include: broad transcriptional alterations and hindered muscle fibre growth^11,23^ (perhaps due to the nuclear disruption which is likely to affect gene expression programmes); myofibril disarray (since desmin and the nuclear envelope are known to contribute to their organisation^24,25^); and altered mechanical properties of muscle fibres (known to be related to the cortical cytoskeleton)^26^. In addition, our results suggest that nuclear and cytoskeletal defects might be a secondary feature and/or source of pathology in other (muscle) diseases, even in those where these structures are not primarily affected.

## Results

### Muscle fibres from NM patients have misshapen and mispositioned nuclei with altered chromatin organisation

To assess myonuclear distribution in patients with NM, single skeletal muscle fibres were teased from biopsy samples, mounted, and stained with rhodamine-labelled phalloidin to visualise actin (thus marking the dimensions of the fibre) and DAPI to label nuclei, and imaged in 3D. Muscle fibres were generally smaller in cross-sectional area (CSA) in patients compared with age-matched control subjects, a common characteristic of NM (as well as some other muscle disorders), although some fibres in *NEB* patients displayed large CSA values, indicating some extent of fibre size disproportion (Fig 1A, B). In one *ACTA1* patient and all three *NEB* patients, there was a higher abundance of myonuclei within muscle fibres compared to control subjects, as observed in nearest neighbour distances (Fig 1C; lower values indicating more densely packed nuclei), and in numbers of nuclei per length of fibre (Fig 1E-H; across a range of fibre sizes/CSAs). In addition, a parameter known as order score was calculated, which is a measure of the regularity of nuclear spacing^27^. Mean order score was significantly lower in all patients compared with control subjects, indicating that nuclei were more unevenly distributed than in controls (Fig 1D). No correlation between fibre CSA and order score was noted in most controls and patients, indicating that nuclear distribution was not noticeably affected by fibre size (Fig S1). Central nuclei are a hallmark of certain neuromuscular disorders including muscular dystrophies; however, this was rare in NM patient fibres, with the vast majority of nuclei being found at the fibre periphery, as expected. Overall, these results suggest that myonuclear positioning is altered in NM patients, and that there is often a higher density of myonuclei within fibres.

**Fig 1.**
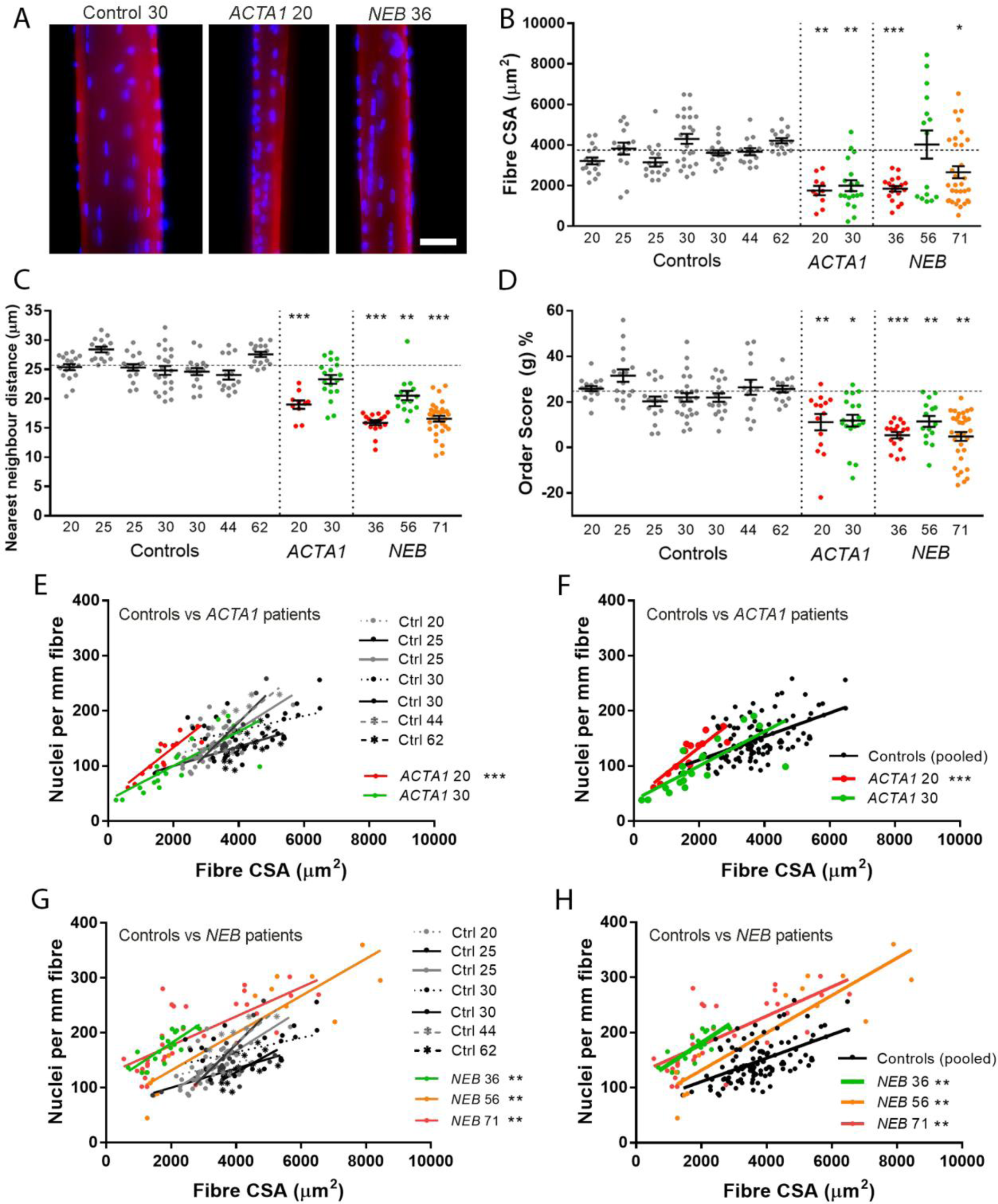
Myonuclei are more abundant and unevenly spaced in patients with nemaline myopathy. Healthy control subjects and patients are denoted with their mutation and age. **(A)** Representative single skeletal muscle fibres from a control subject and patients with *ACTA1* or *NEB* mutations, stained for actin (rhodamine phalloidin; red) and nuclei (DAPI; blue). **(B)** Fibre cross-sectional area (CSA) for controls and patients. **(C)** Nearest neighbour distance between myonuclei within fibres; a smaller distance indicates greater density of nuclei. **(D)** Order score (g), an algorithm to assess the regularity of nuclear spacing^27^; A lower score indicates more irregular spacing and more nuclear clustering. **(E)** Comparison between controls and *ACTA1* patients: relationship between number of nuclei per mm of fibre length, and their CSA. **(F)** As for panel E, but controls are pooled with a single linear regression line, for visual clarity. **(G, H)** As for panels E and F respectively, but comparing controls and *NEB* patients. Individual data points represent an individual skeletal muscle fibre, with mean +/- SEM, or linear regression lines. For column graphs, significance was determined using one-way ANOVA comparing each patient with the closest age-matched controls (*ACTA1* 20 was compared with controls aged 20-25; *ACTA1* 30 and *NEB* 36 was compared with controls aged 30; *NEB* 56 and 71 were compared with controls aged 44 and 62). ANCOVA was used to compare elevations/intercepts and slopes of regression lines. Asterisks refer to the lowest significance level (i.e. highest p value) found for each inter-subject comparison. Scale bar: 50μm. * (P<0.05), ** (P<0.01), *** (P<0.001).

Given that the distribution of nuclei was perturbed in NM patient muscle fibres, we next wanted to determine if there were any alterations within the nuclei themselves. In control subjects, nuclei generally possessed an elliptical shape, and nuclear envelope proteins nesprin-1 and lamin A localised in an even rim around the nuclear edges, as expected (Fig 2A, control subject). However, in two *ACTA1* and two *NEB* patients, >50% of the myonuclei were altered in each case, displaying various features including shape irregularities (Fig 2B), faint staining for nuclear envelope proteins (Fig 2C), nuclear envelope staining resembling bands across the nuclear surface (Fig 2D). Varying proportions of these features were observed in the four patients analysed (Fig 2E). Nuclear shape/elongation was further analysed using area (in the 2D X-Y plane), aspect ratio and circularity measures (Fig 2F-H). Although the nuclear area was on average similar in controls and patients, the variability was significantly greater in patients (Fig 2F). Nuclei in *ACTA1* patients shifted towards a more elongated shape (larger aspect ratio, lower circularity), and nuclei in *NEB* patients towards a more circular shape (lower aspect ratio, higher circularity; Fig 2G, H). Similar nuclear defects are observed in laminopathies, which are primary genetic disorders of the nuclear envelope, and include muscular dystrophies and multisystem disorders caused by mutations in lamin A/C, nesprins and other genes^28^. Our results indicate that severe defects in nuclear envelope and morphology can also occur in diseases caused by muscle contractile dysfunction.

**Fig 2.**
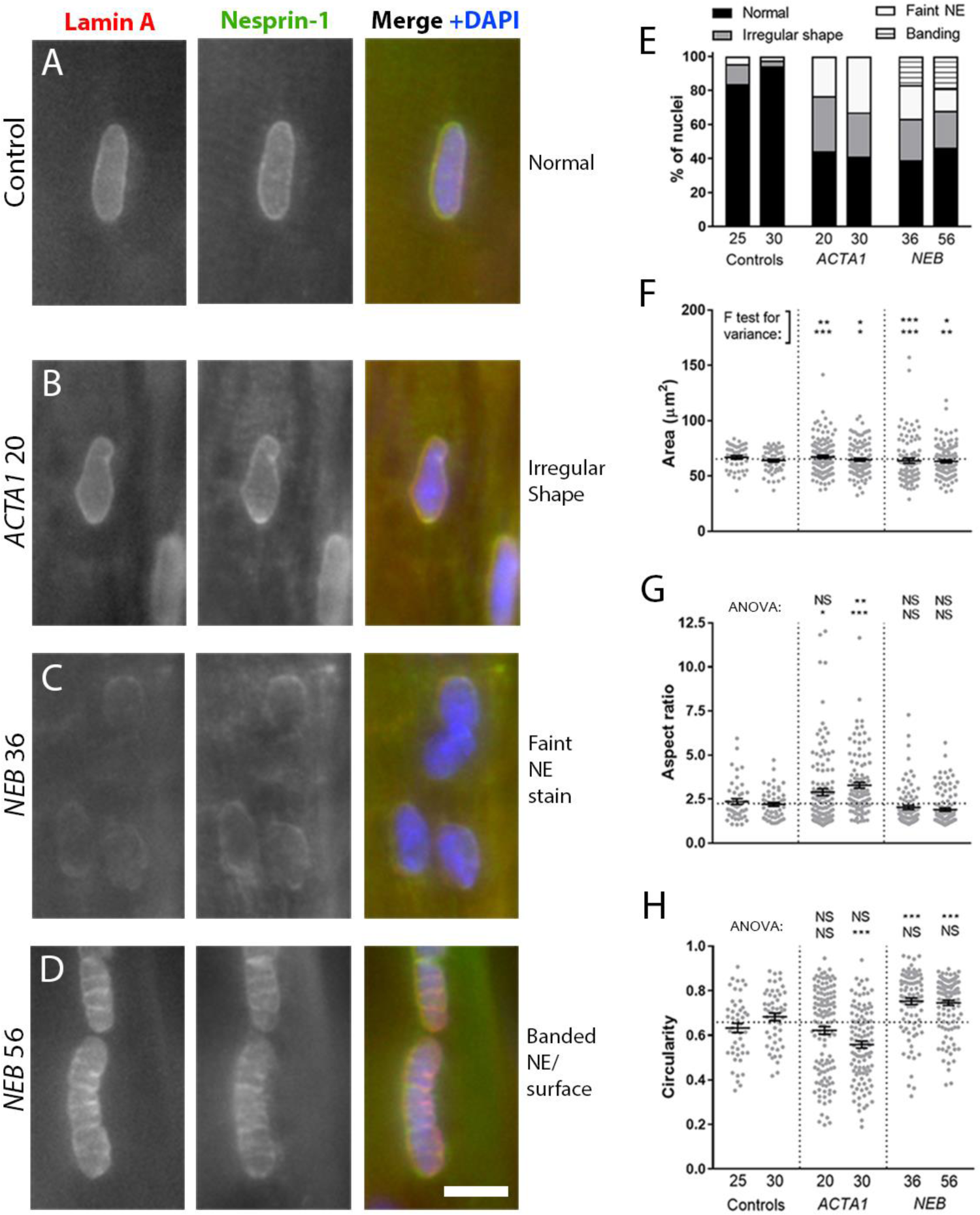
Nuclear morphology and nuclear envelope are altered in skeletal muscle fibres of nemaline myopathy patients. Healthy control subjects and patients are denoted with their mutation and age. **(A-D)** Representative images of myonuclei from healthy controls and NM patients with *ACTA1* or *NEB* mutations, stained for lamin A (left panels), nesprin-1 (middle panels) and merged with DAPI (Lamin A, green; nesprin-1, red; DAPI blue). Myonuclei displaying **(A)** normal shape and nuclear envelope; **(B)** irregular shape; **(C)** faint staining for lamin A and nesprin-1; **(D)** banded pattern across surface. Graph showing proportions of each nuclear phenotype for patients and two controls **(E)**. Graphs showing shape quantifications for nuclei: area **(F),** aspect ratio **(G)** and circularity **(H)**. A larger aspect ratio denotes a more elongated shape; a higher circularity denotes a more circular shape. For shape analysis, only nuclei positioned *en face* were analysed, and those within dense clusters were excluded. Graphs **F - H** show one data point per nucleus analysed, taken from >10 muscle fibres per subject, with mean +/- SEM. Notations for significance denote whether a significant difference is found versus control 25 (upper row) or control 30 (lower row). For graph **F**, an F-test for variance was used, to report differences in spread/variability; for graphs **G** and **H**, ANOVA was used to report differences in mean. Note differences in overall data distribution as well as means. Scale bar: 10μm. NS = not significant * (P<0.05), ** (P<0.01), *** (P<0.001).

Next, ultrastructural imaging was carried out, revealing nuclear morphological and envelope defects in more detail (Fig 3). In NM patients, a proportion of myonuclei resembled those seen in healthy muscle tissue, being of normal shape, having a regular distribution of heterochromatin largely at the nuclear periphery, and with a nuclear envelope consisting of a continuous double membrane (Fig 3A, A’). However, various defects were also noted: clusters of nuclei (Fig 3B); low chromatin density (i.e. high levels of euchromatin; Fig 3C); high chromatin density (i.e. high levels of heterochromatin; Fig 3D, E, G, H); invaginations (marked ‘inv’, Fig 3D); regions of separation between inner and outer nuclear membranes (marked ‘sep’, Fig 3D); discontinuities in the nuclear envelope (arrows, Fig 3E); vacuolations that are continuous with the nuclear envelope (marked ‘v’, Fig 3F-H). Interestingly, in the latter cases, chromatin was never observed to fill the vacuolated regions. Table S3 gives a semi-quantitative analysis of the patients studied, listing the various abnormal features and the relative frequency of occurrence. Fluorescence microscopy of acetylhistone H3 (Lys9/Lys14), a marker of transcriptionally active regions of DNA^29–31^, provided further evidence of chromatin redistribution within nuclei of patients relative to control subjects (Fig 3I-K; image quantifications in Fig S2). Similar observations of nuclear membrane separation and vacuolation have been noted in patients with mutations in the gene encoding nesprin-1^32^, and invaginations and variations in chromatin density have been commonly observed in laminopathies^33,34,35^. Such alterations might be expected to affect nuclear function.

**Fig 3.**
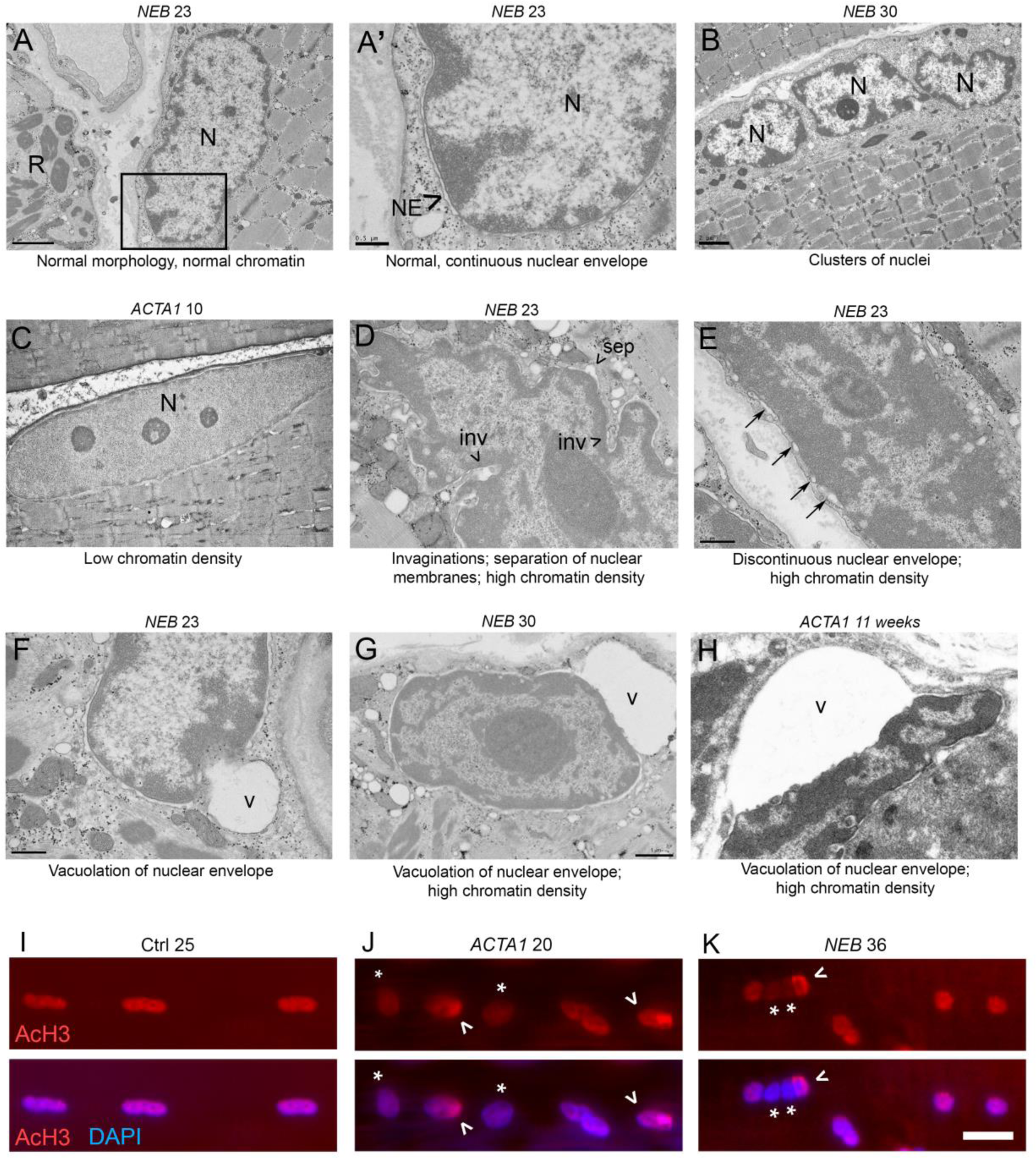
Nuclear envelope and chromatin alterations in myonuclei of nemaline myopathy patients. Patients are denoted with their mutation and age (years unless otherwise specified). **(A, A’)** An example of an apparently relatively normal nucleus (N), with normal morphology, chromatin distribution, and a nuclear envelope consisting of a continuous double membrane (A’ is a magnified image of the boxed region in panel A). Nemaline rods (R) are seen in an adjacent fibre **(A)**. The following nuclear abnormalities were also consistently observed in patients: clusters of nuclei **(B)**; reduced chromatin density **(C)**; increased chromatin density **(D, E, G, H)**; invaginations (**D,** denoted “inv”); separation of the two nuclear membranes (**D,** denoted “sep”); discontinuous patches of nuclear envelope (**E,** arrows); vacuolation, continuous with the nuclear envelope (**F-H**, denoted “v”). **(I-K)** Skeletal muscle fibres were stained with antibody against acetylhistone H3 (Lys9/Lys14), a marker of transcriptionally active regions of DNA (red). DNA is stained with DAPI (blue). Myonuclei from control subjects showed an even distribution of staining throughout the nuclei; however, in patients, a proportion of nuclei showed perturbations in immunolabelling. These included: reduced staining intensity, possibly indicative of global transcriptional downregulation within certain nuclei (asterisks); and uneven distribution of staining, suggesting irregular packing of active genomic regions (arrowheads). Similar results were observed in one other control subject, and one other *ACTA1* and *NEB* patient. 50+ nuclei were observed per subject across ~9 fibres. Quantifications of acetylhistone H3 staining are found in Figure S1. Scale bar (I-K): 20μm.

### Lack of cellular force production is responsible for myonuclear alterations in NM

To investigate potential mechanisms behind aberrant nuclear spacing and morphology in NM, we employed a severe model of the disease, the *Acta1*^H40Y^ mouse^4,36,37^. In a recently published study, a gene therapy approach was used to enhance myosin force output, via the delivery of a myosin light chain isoform (MyL4) that is normally only expressed in developing skeletal muscles (schematic, Fig 4A). This resulted in a partial rescue of the *Acta1*^H40Y^ phenotype, including an improvement in muscle force production and an increase in muscle fibre size^23^. Thus, we aimed to determine whether an increase in force output could also rescue the nuclear defects observed in NM. Like patients with *ACTA1* or *NEB* mutations, skeletal muscle fibres of *Acta1*^H40Y^ mice showed an irregular distribution of myonuclei (Fig 4B), which was quantifiable using the order score parameter (Fig 4C). However, delivery of the *Myl4* transgene resulted in the full restoration of nuclear spacing defects to wild type levels (Fig 4C). In addition, nuclear shape alterations, including increased area and aspect ratio and reduced circularity (Fig 4D-F), were also restored to wild type values in muscles of *Acta1*^H40Y^ mice treated with the transgene. These results suggest that in NM, force impairment results in nuclear spacing and morphological alterations, and that enhancing force production restores these parameters.

**Fig 4.**
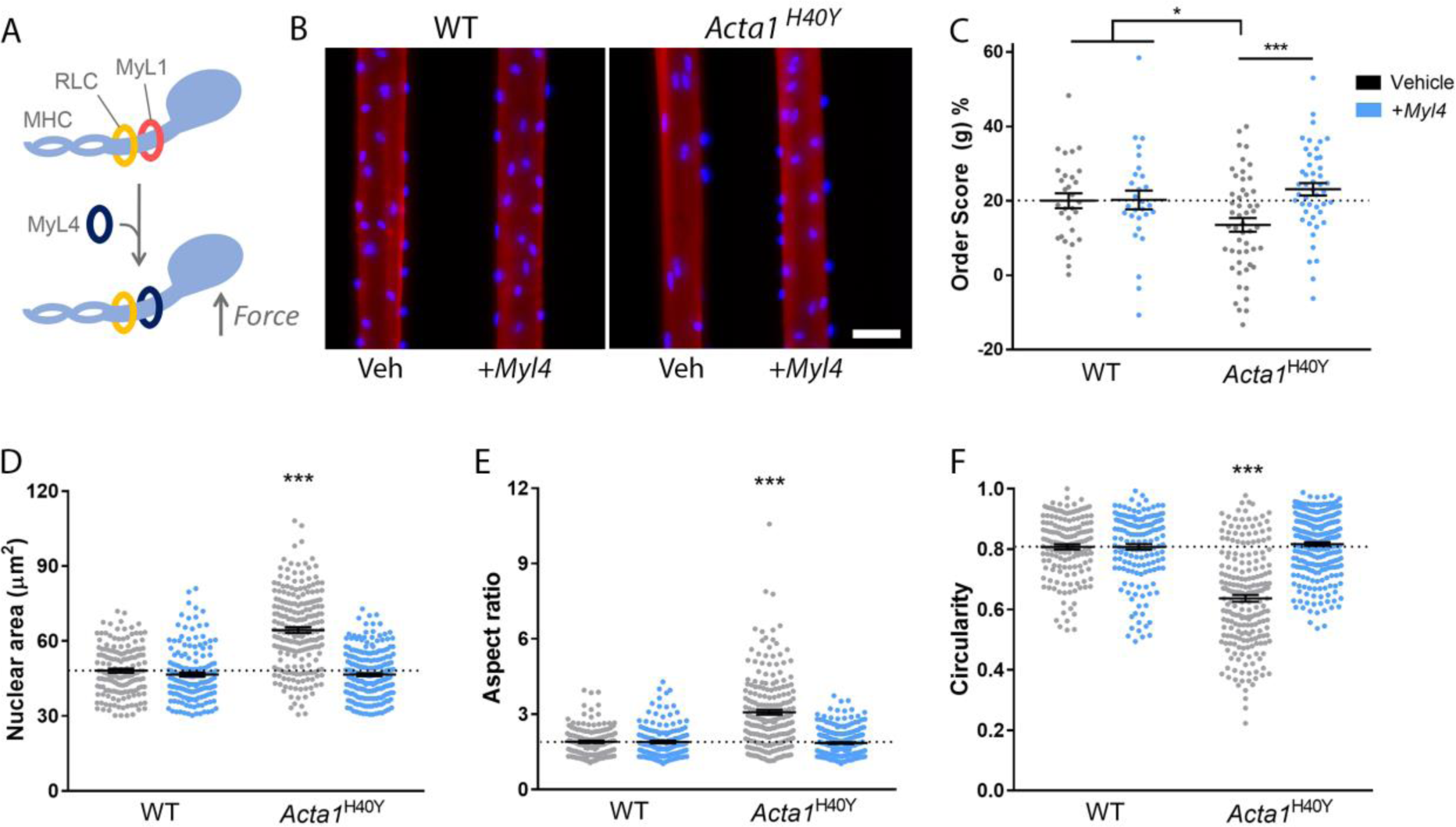
Partial rescue of force production results in full restoration of nuclear spacing and morphology in the *Acta1*^H40Y^ model of nemaline myopathy. **(A)** Scheme for mouse model published previously, where a gene therapy approach was used to deliver a transgene (*Myl4*) into tibialis anterior muscles. *Myl4* encodes a myosin light chain isoform (MyL4) normally only expressed in developing muscles, but when incorporated into the adult myosin complex, results in a myosin with increased force production (MHC, myosin heavy chain; RLC, regulatory light chain; MyL1, the endogenous myosin light chain). **(B)** Representative single skeletal muscle fibres from wild type (WT) and *Acta1*^H40Y^ mice, treated with empty vector (Veh), or *Myl4* transgene. Fibres were stained for actin (rhodamine phalloidin; red) and nuclei (DAPI; blue). **(C)** Order score (g), an algorithm to assess the regularity of nuclear spacing^27^; A lower score indicates more irregular spacing and more nuclear clustering. Nuclear shape measurements: nuclear area as observed in standard x,y planes **(D)**, aspect ratio **(E)** and circularity **(F)**. Note in **C-F** that the alterations in nuclear spacing and morphology observed in mutants were restored to wild type levels when treated with *Myl4* transgene. **(C)** One data point per muscle fibre; **(D-F)** one data point per nucleus analysed, mean +/- SEM, with asterisks denoting significance versus WT/vehicle. Scale bar: 50μm. * (P<0.05), ** (P<0.01), *** (P<0.001).

### Muscle fibres from NM patients and mice have disrupted cytoskeleton networks

In studies using cultured myotubes (an *in vitro* analogue of muscle fibre formation) and in developing *Drosophila*, the lateral spacing/positioning of myonuclei is known to be dependent on microtubules and motor proteins including kinesins and dyneins^13–15,24^. In addition, these motor proteins regulate nuclear shape during their translocation across nascent *Drosophila* fibres^15^. Given that we observe defects in nuclear morphology and spacing in NM patients, we sought to determine whether the microtubule network and/or its associated proteins were perturbed. Microtubules are sensitive to temperature and rapidly depolymerise when tissue is kept cold or frozen^38^, and therefore preservation of their structure is not compatible with routine human muscle biopsy preparation. In lieu of this, we aimed to analyse the localisation of pericentrin, a key microtubule organising centre protein (MTOC) that in skeletal muscle is found at the perinuclear regions, consistent with the role of the myonucleus as a nucleator of microtubules. We found that the localisation of pericentrin was markedly altered in NM patient myonuclei (Fig 5A-D; note the rim of pericentrin around the nuclear surface, as well as nearby bright puncta in control subjects; note the increased accumulation of pericentrin staining at the nuclear surface in patients).

**Fig 5.**
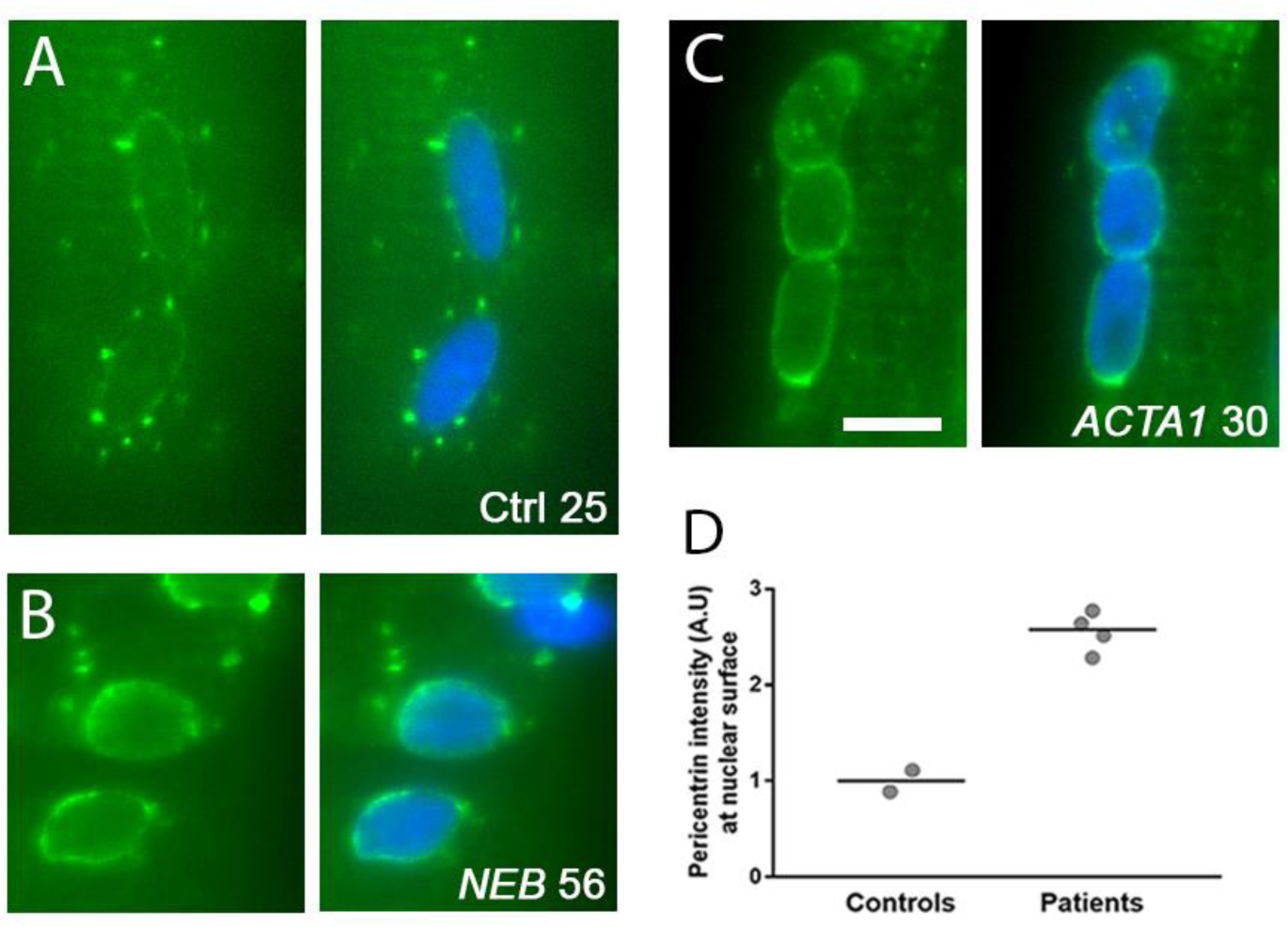
Distribution of the microtubule-organising centre around myonuclei of nemaline myopathy patients. Healthy control subjects and patients are denoted with their mutation and age. **(A-C)** Representative images of nuclei stained for the pericentrin (green) and DAPI (blue). Note specific arrangement of pericentrin in controls (nuclear surface and puncta) versus patients (marked accumulation at the nuclear surface). **(D)** Quantification of fluorescence intensity of pericentrin at the nuclear surface (within 2 μm of nearest nuclear (DAPI)-stained pixel). Scale bar: 10μm.

To further investigate microtubule organisation, we utilised a mouse model of NM with a conditional knockout in the nebulin gene (*Neb* cKO)^6^, and cardiac perfusion fixed with PFA, to allow the proper preservation of microtubule structure. We found that microtubules at the cortex of skeletal muscle fibres were heavily disorganised compared to their control littermates (Fig 6A, B). Note that the microtubules in control animals appear as a regular grid-like lattice, with denser accumulations around the nuclei (arrowheads). In mutants, however, the microtubule network was markedly disorganised, and the extent of accumulation around many of the myonuclei was visibly reduced (asterisks). Further analysis indicated that the microtubule network was denser in the mutant animals (Fig 6E), and that their directionality was disturbed (Fig 6F; a measurement made using the TeDT algorithm tool developed by Liu and Ralston^39^).

**Fig 6.**
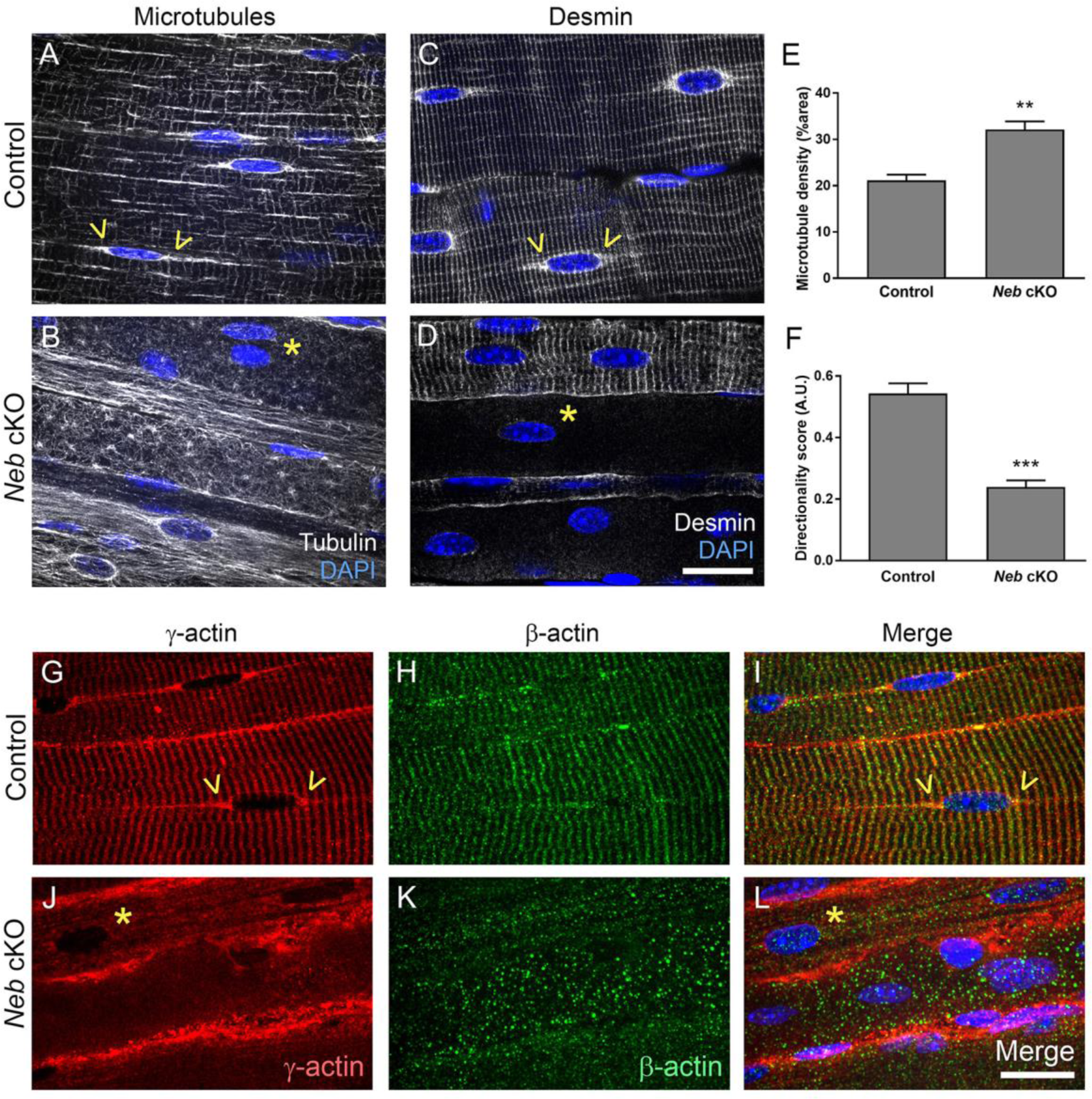
Severe disruption of the cortical cytoskeleton in the *Neb* cKO mouse model of nemaline myopathy. Representative confocal micrographs of skeletal muscle fibres in whole-mount extensor digitorum longus muscles of control **(A, C)** and *Neb* cKO mice **(B, D)**; tissue was immunolabelled with antibodies to β-tubulin **(A, B)** or desmin **(C, D)** and nuclei were stained with DAPI (blue). In control muscles, the skeletal muscle fibre cortex shows a regular grid-like pattern for microtubules and a striated appearance for desmin; both are largely disorganised or reduced in mutants. Note the clustering of microtubules and desmin at the periphery of nuclei, often altered or lacking in mutants. Quantifications of microtubule density **(E)**, and directionality using previously developed algorithms^39^ **(F)**. Cytoplasmic actin stains for control **(G-I)** and Neb cKO **(J-K)** mice; γ-actin **(G, J)**, β-actin **(H, K)** and merged images **(I, L)**. Note that microtubules, desmin and γ-actin show specific accumulation at the surface and poles of nuclei (arrowheads), which is frequently reduced or missing in mutants (asterisks). Observations were similar in 3 mutants and 3 control littermates (comprising 1 wild type and 2 heterozygotes which are unaffected). Graphs, mean +/- SEM. Scale bars: 20μm. * (P<0.05), ** (P<0.01), *** (P<0.001).

Desmin is a muscle-specific intermediate filament protein that couples myofibrils at the Z-discs, and links to other organelles including nuclei. We found that in most (~90%) of Neb cKO fibres, the normal Z-disc localisation of desmin was absent, and that its accumulation at the nuclear surface was not discernible (asterisks, Fig 6C, D). Similar results were observed for cytoplasmic actins (β- and γ-) which form part of the cortical cytoskeleton. In control animals, both β- and γ-actin localised in striations, previously shown to be in alignment with Z-discs of the sarcomere (Fig 6G-I)^40–42^. However, in *Neb* cKO animals, both the striations and the nuclear regions of cytoplasmic actins were virtually absent, although γ-actin appeared as accumulations at the fibre periphery (Fig 6J-L). Together these results indicate that microtubule, desmin and cytoplasmic actin networks are markedly altered in this mouse model of NM.

### Cytoskeletal organisation defines myonuclear properties

To investigate whether disorganisation of the cytoskeleton might cause nuclear abnormalities, intact muscle fibres were isolated enzymatically from mouse extensor digitorum longus (EDL) muscles, and cultured with and without drugs that modulate microtubule structure and dynamics. Overnight treatment with nocodazole resulted in an almost complete removal of microtubules (Fig 7A, B, E), whereas treatment with taxol or Epothilone D (EpoD) resulted in increased microtubule density and reduced directionality score compared with control/vehicle treated fibres (Fig 7C-F). Nocodazole resulted in small shifts in nuclear morphology towards a more elongated, less circular phenotype (Fig 7H, I; arrowhead in Fig 7B indicates an example of an abnormally elongated nucleus), although nuclear area was not affected (Fig 7G). Treatment with taxol or EpoD resulted in an increase in nuclear area (X-Y planes) compared with control fibres (Fig 7J, top row of panels; Fig 7K), although no changes in aspect ratio or circularity were found (Fig 7M, N). In addition, using 3D Z-stacks, we found similar nuclear volumes between control, taxol- and EpoD-treated fibres (Fig 7L). This suggests that the increases in nuclear area observed with taxol and EpoD are caused by the nuclei spreading out/becoming flatter, rather than by an overall expansion of their total volume. Consistent with this, the lower panels in Fig 7J show examples of the Z plane of the Z-stacks, with nuclei appearing flatter in this dimension. These results imply that the microtubule network exerts tension on the nuclear surface to regulate nuclear flattening.

**Fig 7.**
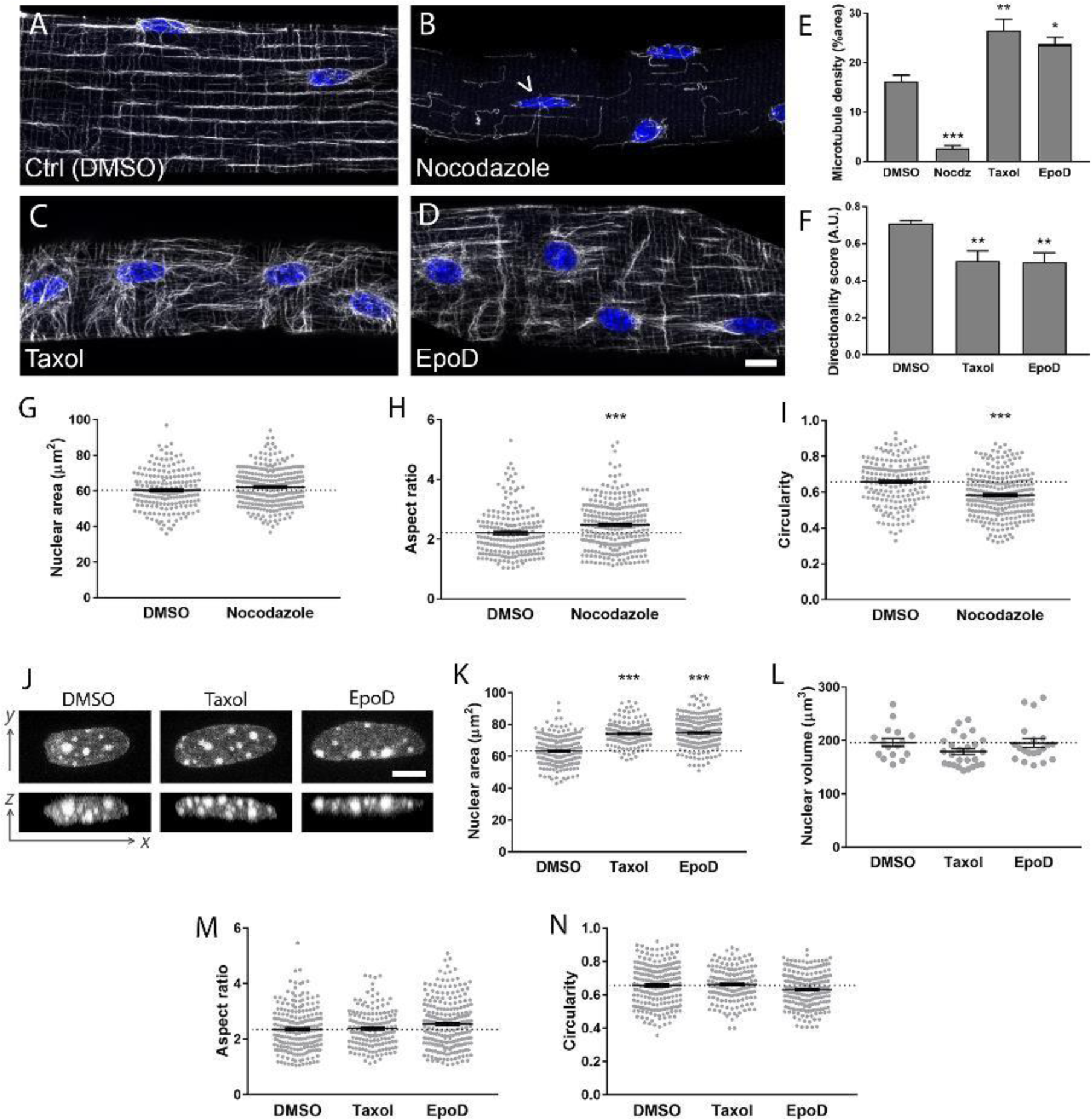
Pharmacological disruption of microtubule network structure results in alterations to nuclear morphology. Enzymatically dissociated single muscle fibres from wild type mouse extensor digitorum muscles treated overnight with **(A)** DMSO (control), **(B)** Nocodazole, **(C)** taxol and **(D)** Epothilone D (EpoD). Microtubules are shown in grey (anti-β-tubulin), and nuclei in blue (DAPI). Quantifications of microtubule density **(E)**, and directionality using previously developed algorithms^39^ **(F)**, in drug-treated fibres. Nuclear shape measurements for control versus nocodazole treated fibres: nuclear area as observed in 2 dimensions **(G)**, aspect ratio **(H)** and circularity **(I)**; these results show a modest shift towards more elongated nuclei with nocodazole treatment (arrowhead in B shows example of an elongated nucleus). Representative images of nuclei from fibres treated with DMSO, taxol and EpoD **(J)**: upper row, nuclei as observed in standard x,y planes; lower row, orthogonal views of nuclei as seen in x,z planes. Nuclear shape measurements for control versus taxol and EpoD treated fibres: nuclear area as observed in standard x,y planes **(K)**, nuclear volume **(L),** aspect ratio **(M)** and circularity **(N)**. All graphs show mean +/- SEM; one data point per nucleus in panels **G-I** and **K-N**. Microtubule and nuclear measurements were taken from multiple regions in 8 fibres per condition, spread across 2-3 separate experiments for each condition. Asterisks denote significance versus DMSO/control treated fibres. Scale bars: 10μm **(A-D)** and 5μm **(J)**. * (P<0.05), ** (P<0.01), *** (P<0.001).

It should be noted that myonuclear spacing was preserved in all drug-treated fibres relative to control, even after 72 hours treatment (Fig 7A-D and Fig S3), suggesting that at least in non-contracting isolated muscle fibres, microtubule disruption has no major impact on nuclear movement. Also, there were no gross changes in nesprin-1 localisation with any of the drug treatments, suggesting that microtubule disruption did not have any major effects on the nuclear envelope (Fig S3).

## Discussion

In this study, we identify a range of nuclear and cytoskeletal defects that occur as signs of *ACTA1* or *NEB*-related NM pathology. We show that defects in nuclear morphology and spacing are caused by the impairment in force production that is a characteristic of the disease. We also demonstrate the role of the microtubule cytoskeleton in the regulation of nuclear shape.

Myonuclear spacing defects (Fig 1) have also been observed in other mutant mouse models, including those in genes encoding nuclear envelope proteins (SUN1/SUN2 double KO, Nesprin-1 KO^43–45^), and also in ageing^10^. Interestingly, this effect appears to be specific, since knockout of other nuclear envelope proteins such as SUN1, SUN2 or nesprin-2 does not significantly alter myonuclear organisation^31,43,44,46^. Currently there is little insight into whether defects in nuclear spacing contribute to cellular pathology, or whether they are a minor consequence of other events. However, given that regular spacing of myonuclei is a highly conserved feature across taxa, it is assumed that it is important for proper muscle fibre function, such as inter-nuclear cooperation and the efficient distribution of gene products throughout the cell^11^. Therefore, one might envisage that any deviation from a “normal” nuclear arrangement would result in sub-optimal muscle fibre function.

Studies to ascertain the mechanisms of nuclear spacing in skeletal muscle have largely taken place in myotubes and embryos of *Drosophila* and mouse, and have identified a number of mediators including nesprins, microtubules, MTOC proteins and the motor proteins kinesin and dynein^12,15,45,47^. Myotubes, as an *in vitro* system are analogous to the events during embryonic development, whereby myonuclei are buried in the centre/core of the fibre, where their spacing takes place along the axis of the fibre. As such, these cells are not anatomically or developmentally equivalent to mature fibres, in which nuclei are anchored at the fibre periphery. In these examples of NM, it is not clear whether the nuclear organisation defects are a result of aberrant spacing occurring during development, or in events that occur in maturity. However, our results do suggest that nuclear organisation can be remodelled in mature fibres, since delivery of the *Myl4* transgene into adult *Acta1*^H40Y^ mice resulted in nuclear spacing that was restored to that of wild type levels (Fig 4). One of the key features of muscle tissue in NM (as well as some other disorders) is a shift towards type I fibres. This alone is unlikely to explain the alterations that we observed in NM patients, since no clear difference in the regularity of nuclear spacing was observed across human fibre types^48^.

Nuclear envelope and shape defects are also a feature of NM patients (Fig 2-3), which more typically is associated with disorders caused by mutations affecting nuclear envelope proteins. Indeed, mutations in the gene encoding nesprin-1 result in areas of separation and vacuolation of the nuclear envelope (similar in appearance to our ultrastructural findings)^32^, and shape alterations and invaginations have been observed in various laminopathy cases^28,33,49^. What is less clear, however, is to what extent nuclear abnormalities are present in other neuromuscular disorders, although it has often been noted as a secondary feature of some myopathies^50,51^ (personal communication, Prof Caroline Sewry), as well as in cultured fibroblasts from patients with Duchenne muscular dystrophy^52^. Given that defects in the nuclear envelope are the primary cause of a spectrum of diseases, it is likely that the secondary defects observed in NM have pathological consequences. Perhaps associated with this is the frequent observation of chromatin abnormalities in NM patients by EM and light microscopy (Fig 3, Supplementary Fig S2), since both nuclear shape and the nuclear envelope are known to play a role in transcriptional regulation via chemical and mechanical control of chromatin organisation^17–22^. Analogously, similar effects on chromatin are seen in myonuclei of patients with mutations affecting nesprins or lamins^33,34,35^. Indeed, broad alterations in the transcriptional profile of skeletal muscle is observed in patients with NM (including in genes related to metabolism and calcium homeostasis), and this may partially be the result of nuclear shape and envelope alterations and/or reorganised chromatin^53^.

In the delivery of the *Myl4* transgene to adult *Acta1*^H40Y^ mouse muscles, contractile force is augmented^23^ and nuclear spacing and morphology is restored to wild type levels (Fig 4). This suggests that (i) a lack of force originating at the sarcomere is responsible for these nuclear defects, and that (ii) this can be reversed by an increase in contractile capacity. The mediators of this effect are unclear, but may involve direct force transmission to the nuclei via e.g. microtubule, actin or desmin cytoskeletons, or interactions through second messengers that may be responsive to mechanical input. It cannot be excluded that myosin light chain 4 itself is acting as the key mediator of this effect, rather than force transmission per se; however, this is unlikely, as many mechanosensitive molecules are known to be responsive to force transmission^54–56^, and any of these could be responsible for changes in nuclear shape and/or distribution. Overall, these results raise the possibility that nuclear morphology and spacing alterations might not be specific to NM, since various other classes of myopathy are also associated with impaired contraction^1^.

Another key finding was cytoskeletal defects in the *Neb* cKO model NM, involving microtubules, desmin and non-sarcomeric actins (Fig 5-6). Microtubules also show increased density and disorganisation in dystrophic mice with dystrophin or sarcoglycan deficiency, or mice with MAP6 ablation^57,58^. In skeletal muscle, microtubules are known to have roles in modulating fibre stiffness and contraction, and in signalling via reactive oxygen species (ROS)^26^. Indeed, the increased microtubule density in dystrophic mice results in elevated ROS, over-activation of stretch-sensitive Ca^2+^ channels, and worsened pathology^59^. Defects in non-sarcomeric actins and desmin were striking in that the vast majority of fibres showed markedly reduced/virtually absent localisation of all of these components. Ablation of either of these actins specifically in skeletal muscle causes mild progressive myopathies^60^, and loss of desmin causes disruptions of muscle architecture^25^. Intriguingly, myofibrils are frequently misaligned and disordered in NM^8^, and this might be due to (i) the loss of desmin, and/or (ii) the defects at the nuclear envelope, since both have been shown to be involved in the proper arrangement of myofibrils^24,25^. One key role of desmin and actins is the lateral transmission of force to the fibre periphery, where actin and microtubules bind to the dystrophin-associated glycoprotein complex at the plasma membrane^41,42,57^. Therefore, the mislocalisation of these components is likely to have implications for the mechanical properties of the fibre. Cytoskeletal components also have important roles in anchorage at the nucleus, and the normal localisation of microtubules, γ-actin and desmin at the nuclear surface was also largely reduced in *Neb* cKO mice. The cytoskeleton is likely to transmit strain from the mechanical forces of contraction to the nuclei, which may be important for fibre integrity and/or regulation of gene expression^61^. Consistent with this, disruption of microtubules with several agents resulted in alterations to nuclear shape (Fig 7). Although microtubules are known to regulate nuclear spacing during development, no changes in nuclear distribution were observed when fibres were treated with nocodazole, taxol or EpoD (Fig 7, Fig S2). Possible explanations for this include: (i) other cytoskeletal systems are instead responsible for nuclear spacing in mature fibres, or are able to compensate when microtubules are disrupted; (ii) nuclei are more mobile in actively contracting fibres, which was not the case in these experiments.

The range of disease severity varies greatly in NM, and death in childhood is frequent. In this study, the patients were almost entirely of adult age, representing the milder end of the spectrum. This was an experimental design choice, due to the availability of healthy human control tissue at adult, but not childhood ages. Therefore, it is impossible to pinpoint whether similar effects occur in typical congenital cases of NM. To date, our understanding of muscle development at the cellular level during infancy and childhood is incomplete, and there is little data on nuclear spacing, morphology or cortical cytoskeleton, although the myonuclear domain is known to steeply increase during the first few years of life^62^. As such, interpretation of cellular organisation in congenital NM patients would be difficult, without appropriate information on the same parameters during healthy development.

In summary, these results demonstrate that skeletal muscle from NM patients and mouse models display defects in the non-sarcomeric cytoskeleton and in nuclear positioning and integrity. They indicate that abnormal nuclear spacing and morphology are the result of the impaired contractile force production that is a key feature of this disease. In addition, this study highlights the role of a properly organised cytoskeleton in the regulation of nuclear morphology. Although nuclear defects are observed in other diseases, including those caused by mutations in nuclear envelope proteins, these findings are somewhat unexpected, given that NM pathology originates at the sarcomere^2,63^. They might explain some of the features observed in NM, such as broad transcriptional changes and hindered muscle fibre growth (possibly related to alterations in nuclear envelope and chromatin organisation, which is likely to affect programmes of gene expression^17,18,22^), myofibrillar disarray (due to the roles of desmin and the nuclear envelope in sarcomere organisation^24,25^) and altered fibre mechanical properties (perhaps related to cytoskeletal arrangement^26^). This study raises the possibility that nuclear and cytoskeletal defects may be an overlooked feature and/or source of pathology in other (muscle) diseases.

## Methods

### Human subjects

All tissue was consented, stored and used in accordance with the Human Tissue Act, UK, under local ethical approval (REC 13/NE/0373). Details of patients for light microscopy and electron microscopy are given in tables S1 and S2 respectively.

### Mouse models

*Acta1*^H40Y^ mouse tibialis anterior muscle samples, with and without transfer of the *Myl4* gene remained frozen from a previous study^23^. To summarise, data were collected from 4 wild type and 4 mutant male mice. At 4 weeks of age, the compartment containing the tibialis anterior muscle was injected with rAAV6 virus containing the *Myl4* transgene, and contralateral legs served as controls, being injected with virus lacking the functional gene. Mice were sacrificed 4 weeks later at 8 weeks of age. Colony maintenance and experiments were approved by the Uppsala Local Ethical Committee on Animal Research.

*Neb* cKO mice^6^ were maintained at the University of Arizona in accordance with the US National Institutes of Health guidelines “Using Animals in Intramural Research”. Mutants and WT/heterozygous littermates were sacrificed at 3 months of age by cardiac perfusion with 4% paraformaldehyde (PFA)/PBS, to properly preserve microtubule structure. Extensor digitorum longus muscles were dissected for whole-mount staining of cytoplasmic actins, desmin or microtubules (see below).

### Antibodies

Primary antibodies were as follows (species, isotype, manufacturer and dilution are given): lamin A (mouse monoclonal IgG3, Abcam, 1:200); nesprin-1 (rabbit monoclonal IgG, Abcam, 1:400); pericentrin (rabbit polyclonal IgG, Abcam, 1:200); β-tubulin (clone TUB2.1, mouse monoclonal IgG1, Santa Cruz, 1:500); desmin (clone D33, mouse monoclonal IgG1, Dako, 1:400); β-actin (rabbit polyclonal IgG, Abcam, 1:300); γ-actin (clone 2A3, mouse monoclonal IgG2b, Bio-Rad, 1:300); acetyl lys9/lys14 histone H3 (rabbit polyclonal IgG, Cell Signaling, 1:200).

### Enzymatic isolation and culture of intact single muscle fibres

Intact single muscle fibres were prepared as described previously, using enzymatic dissociation with collagenase I and gentle trituration (Sigma Aldrich)^64^. After isolation, fibres were plated into 6-well plates (~30 fibres per well) in DMSO/high glucose/GlutaMAX supplement/pyruvate (Thermo Fisher Scientific, Cat# 31966021), containing 10% horse serum and 1% Penicillin/streptomycin solution. Freshly isolated fibres were treated overnight with nocodazole (20 μM), taxol (10 μM) or epothilone D (10 μM). Final DMSO (vehicle) concentration was 0.5% in all cases. To assess myonuclear spacing, fibres were cultured for 72 hours in the presence of the aforementioned drugs.

### Immunohistochemistry (single muscle fibres)

Fibres were fixed in 4% PFA/PBS for 15 mins, and washed 3x in PBS. Fibres were permeabilised in 0.1% triton-X/PBS for 10 mins, washed 3x and blocked in 10% normal goat serum/PBS for 1 hour. Fibres were then treated with primary antibodies in blocking solution overnight (β-tubulin) or for 3 hours (lamin A, nesprin-1, pericentrin) at 4**°**C. Fibres were washed 3x in PBS for a total of 30 mins, and then treated with Alexa 594 or 488-conjugated secondary antibodies and DAPI (all at 1:1000 in PBS) for 3 hours. Fibres were washed 3x in PBS for a total of 30 mins and mounted in Fluoromount mounting medium (Southern Biotech) with coverslip (thickness #1.5).

### Immunohistochemistry (whole mount muscles)

*Neb* cKO mice and WT/heterozygous littermates (all female) were sacrificed at 3 months of age by cardiac perfusion with 4% paraformaldehyde (PFA)/PBS, to properly preserve microtubule structure. Extensor digitorum longus muscles were dissected for whole mount staining of cytoplasmic actins, desmin or microtubules. Muscles were then permeabilised in 0.5% triton-X/PBS (20 mins) and 0.1% Triton-X/PBS (20 mins), with each solution being replaced at least once during the incubation. Samples were then blocked in mouse-on-mouse block/PBS for 3 hours, and then blocked in 8% bovine serum albumin overnight. Primary antibodies (tubulin, cytoplasmic actins) in blocking buffer were applied for 5 hours, followed by 2 hours washing in 0.1% triton-X/PBS.

### Fluorescence Imaging

Fibres were imaged on a Zeiss Axiovert 200 spinning disc confocal microscope equipped with BD CARV II and a motorised Z drive at x20 magnification (for imaging of nuclear morphology and nuclear envelope). For nuclear number and distribution, Z-stacks with 1 μm Z increments were taken through the entire depths of fibres, as described previously^65,66^. For imaging of cytoskeleton and nuclear volume, a Nikon A1 laser scanning confocal microscope with a x100 oil immersion objective (1.4 NA) was used, with Z-stacks taken with 0.3 μm Z increments (Nikon Imaging Centre, King’s College London).

### Image Analysis

Analysis of nuclear number and spacing: Coordinates of myonuclei were identified in 3D within Z-stacks of muscle fibres. A custom-made Matlab programme was used to a measure fibre CSA, nuclear number, nearest neighbour distances and order score (‘g’) of nuclei within fibres, as described previously^27,65^.

Analysis of nuclear shape parameters: For 2- and 3-dimensional measurements (area, aspect ratio, circularity and volume), nuclei in the DAPI channel were thresholded by pixel intensity until fully highlighted. Inbuilt ImageJ functions were used to measure 2D parameters and Voxel Counter plugin for volume. For accurate shape analysis, nuclei positioned on the sides or the backs of fibres (relative to the microscope objective) were excluded.

Microtubule quantifications: density (% area) was calculated on binary converted images in ImageJ. Microtubule directionality was calculated using the TeDT tool^39^.

### Statistics

Graphs were prepared and analysed in Graphpad Prism. Linear regression lines and statistical comparisons were performed using inbuilt algorithms (ANCOVA test was used to compare elevations/intercepts and slopes of different regression lines). For column comparisons, a two-tailed t test was used to compare 2 groups, and a one-way ANOVA with Tukey post-correction was used to compare more than 2 groups. For studies with animals, no significant differences were observed between animals of the same genotype, and as such, individual values were pooled. Asterisks denote the following statistical significance levels: * (P<0.05), ** (P<0.01), *** (P<0.001).

## Acknowledgments

We thank Prof Caroline Sewry for her invaluable knowledge, and guidance with the interpretation of the data and the preparation of the manuscript. We thank Drs Wenhua Liu and Evelyn Ralston (National Institute of Arthritis and Musculoskeletal and Skin Diseases, NIH, Bethesda, MD, USA) for providing the microtubule directionality (TeDT) tool. We also thank the Nikon Imaging Centre at King’s College London for the provision of equipment for, and assistance with confocal imaging. This work was generously funded by the Medical Research Council of the UK.

## Author contributions

J.A.R., J.L., E.C.H. and J.O. contributed to the design the study. All authors contributed to the writing and preparation of the manuscript. J.A.R., Y.L., J.S.K., M.T., M.R., M.M., J.L., N.F., and J.O. carried out experiments. J.A.R., Y.L., M.T., M.H., M.R. and M.M. analysed data. M.M., C.F., H.J., J.V., N.W., E.Z., C.S. and C.W-P. provided patient samples and clinical data. P.S.Z, H.G. and E.C.H. provided animal models.

## References

1. Jungbluth, H. et al. Congenital myopathies: disorders of excitation–contraction coupling and muscle contraction. Nat. Publ. Gr. (2018). doi:10.1038/nrneurol.2017.191

2. Chan, C. et al. Myopathy-inducing mutation H40Y in ACTA1 hampers actin filament structure and function. Biochim. Biophys. Acta - Mol. Basis Dis. 1862, 1453–1458 (2016)).

3. Ochala, J., Ravenscroft, G., Mcnamara, E., Nowak, K. J. & Iwamoto, H. X-ray recordings reveal how a human disease-linked skeletal muscle a -actin mutation leads to contractile dysfunction. J. Struct. Biol. 192, 331–335 (2015)).

4. Lindqvist, J., Cheng, A. J., Renaud, G., Hardeman, E. C. & Ochala, J. Distinct underlying mechanisms of limb and respiratory muscle fiber weaknesses in nemaline myopathy. J. Neuropathol. Exp. Neurol. 72, 472–81 (2013)).

5. Ochala, J. et al. Congenital myopathy-causing tropomyosin mutations induce thin filament dysfunction via distinct physiological mechanisms. Hum. Mol. Genet. 21, 4473–4485 (2012)).

6. Li, F. et al. Nebulin deficiency in adult muscle causes sarcomere defects and muscle-type-dependent changes in trophicity: Novel insights in nemaline myopathy. Hum. Mol. Genet. 24, 5219–5233 (2015)).

7. Gokhin, D. S., Ochala, J., Domenighetti, A. A. & Fowler, V. M. Tropomodulin 1 directly controls thin filament length in both wild-type and tropomodulin 4-deficient skeletal muscle. Development 142, 4351–4362 (2015).

8. Tonino, P. et al. Reduced myofibrillar connectivity and increased Z-disk width in nebulin-deficient skeletal muscle. J. Cell Sci. 123, 384–91 (2010)).

9. Winter, J. M. D. et al. Mutation-specific effects on thin filament length in thin filament myopathy. Ann. Neurol. 79, 959–969 (2016)).

10. Bruusgaard, J. C., Liestøl, K. & Gundersen, K. Distribution of myonuclei and microtubules in live muscle fibers of young, middle-aged, and old mice. J. Appl. Physiol. 100, 2024–2030 (2006)).

11. van der Meer, S. F. T., Jaspers, R. T. & Degens, H. Is the myonuclear domain size fixed? J. Musculoskelet. Neuronal Interact. 11, 286–297 (2011).

12. Gimpel, P. et al. Nesprin-1α-Dependent Microtubule Nucleation from the Nuclear Envelope via Akap450 Is Necessary for Nuclear Positioning in Muscle Cells. Curr. Biol. 27, 2999–3009.e9 (2017).

13. Falcone, S. et al. N-WASP is required for Amphiphysin-2/BIN1-dependent nuclear positioning and triad organization in skeletal muscle and is involved in the pathophysiology of centronuclear myopathy. EMBO Mol. Med. 6, 1455–75 (2014)).

14. Metzger, T. et al. MAP and kinesin-dependent nuclear positioning is required for skeletal muscle function. Nature 484, 120–124 (2012).

15. Folker, E. S., Schulman, V. K. & Baylies, M. K. Translocating myonuclei have distinct leading and lagging edges that require Kinesin and Dynein. Development 141, 355–66 (2014).

16. Roman, William, Martins, Joao P., Carvalho, Filomena A., Voituriez, Raphael, Abella, Jasmine V.G., Santos, Nuno C., Cadot, Bruno, Way, Michael, Gomes, E. R. Myofibril contraction and cross-linking drive nuclear movement to the periphery of skeletal muscle. Nat. Cell Biol. (2017). doi:10.1038/ncb3605

17. Meinke, P. & Schirmer, E. C. LINC’ing form and function at the nuclear envelope. FEBS Lett.589, 2514–2521 (2015).

18. Hetzer, M. W. The Nuclear Envelope. Online 1411, 241–254 (2010).

19. Tajik, A. et al. Transcription upregulation via force-induced direct stretching of chromatin. doi:10.1038/NMAT4729

20. D’Alessandro, M. et al. Amphiphysin 2 Orchestrates Nucleus Positioning and Shape by Linking the Nuclear Envelope to the Actin and Microtubule Cytoskeleton. Dev. Cell 35, 186–198 (2015).

21. Kim, J. K. et al. Nuclear lamin A/C harnesses the perinuclear apical actin cables to protect nuclear morphology. Nat. Commun. 8, 1–13 (2017)).

22. Webster, M., Witkin, K. L. & Cohen-Fix, O. Sizing up the nucleus: nuclear shape, size and nuclear-envelope assembly. J. Cell Sci. 122, 1477–1486 (2009)).

23. Lindqvist, J. et al. Modulating Myosin Restores Muscle Function in a Mouse Model of Nemaline Myopathy. (2016). doi:10.1002/ana.24619

24. Auld, A. L. & Folker, E. S. Nucleus-dependent sarcomere assembly is mediated by the LINC complex. Mol. Biol. Cell 27, 2351–9 (2016).

25. Capetanaki, Y., Milner, D. J. & Weitzer, G. Desmin in muscle formation and maintenance: knockouts and consequences. Cell Struct. Funct. 22, 103–116 (1997)).

26. Kerr, J. P. et al. Detyrosinated microtubules modulate mechanotransduction in heart and skeletal muscle. Nat. Commun. 6, 8526 (2015).

27. Bruusgaard, J. C., Liestøl, K., Ekmark, M., Kollstad, K. & Gundersen, K. Number and spatial distribution of nuclei in the muscle fibres of normal mice studied in vivo. J. Physiol. 551, 467– 78 (2003).

28. Janin, A. & Gache, V. Nesprins and lamins in health and diseases of cardiac and skeletal muscles. Front. Physiol. 9, 1–12 (2018)).

29. Kirby, T. J. et al. Myonuclear transcription is responsive to mechanical load and DNA content but uncoupled from cell size during hypertrophy. Mol. Biol. Cell 27, 788–798 (2016).

30. Ravel-Chapuis, A., Vandromme, M., Thomas, J. L. & Schaeffer, L. Postsynaptic chromatin is under neural control at the neuromuscular junction. EMBO J. 26, 1117–1128 (2007)).

31. Gnocchi, V. F. et al. Uncoordinated transcription and compromised muscle function in the Lmna-null mouse model of Emery-Dreifuss muscular dystrophy. PLoS One 6, 1–12 (2011).

32. Kolbel, H. et al. Characteristic clinical and ultrastructural findings in nesprinopathies. Eur. J. Paediatr. Neurol. 23, 1–8 (2018)).

33. Baumann, M. et al. Homozygous SYNE1 mutation causes congenital onset of muscular weakness with distal arthrogryposis: a genotype-phenotype correlation. Eur. J. Hum. Genet. 262–266 (2016). doi:10.1038/ejhg.2016.144

34. Fidziańska, A. & Hausmanowa-Petrusewicz, I. Architectural abnormalities in muscle nuclei. Ultrastructural differences between X-linked and autosomal dominant forms of EDMD. J. Neurol. Sci. 210, 47–51 (2003)).

35. Sewry, C. A. et al. Skeletal muscle pathology in autosomal dominant Emery-Dreifuss muscular dystrophy with lamin A/C mutations. Neuropathol. Appl. Neurobiol. 27, 281–290 (2001)).

36. Nguyen, M. A. T. et al. Hypertrophy and dietary tyrosine ameliorate the phenotypes of a mouse model of severe nemaline myopathy. Brain 134, 3513–3526 (2011).

37. Tinklenberg, J. et al. Treatment with ActRIIB-mFc Produces Myofiber Growth and Improves Lifespan in the Acta1 H40Y Murine Model of Nemaline Myopathy. Am. J. Pathol. 186, 1568– 1581 (2016).

38. Wallin, M. & Strömberg, E. Cold-stable and cold-adapted microtubules. Int. Rev. Cytol. 157, 1– 31 (1995).

39. Liu, W. & Ralston, E. A new directionality tool for assessing microtubule pattern alterations. Cytoskeleton 71, 230–240 (2014).

40. Rybakova, I. N., Patel, J. R. & Ervasti, J. M. The dystrophin complex forms a mechanically strong link between the sarcolemma and costameric actin. J. Cell Biol. 150, 1209–1214 (2000)).

41. Prins, K. W., Call, J. a, Lowe, D. a & Ervasti, J. M. Quadriceps myopathy caused by skeletal muscle-specific ablation of β(cyto)-actin. J. Cell Sci. 124, 951–7 (2011)).

42. Sonnemann, K. J. et al. Cytoplasmic gamma-actin is not required for skeletal muscle development but its absence leads to a progressive myopathy. Dev. Cell 11, 387–97 (2006).

43. Zhang, X. et al. Syne-1 and Syne-2 play crucial roles in myonuclear anchorage and motor neuron innervation. Development 134, 901–908 (2007).

44. Lei, K. et al. SUN1 and SUN2 play critical but partially redundant roles in anchoring nuclei in skeletal muscle cells in mice. Proc. Natl. Acad. Sci. U. S. A. 106, 10207–10212 (2009)).

45. Stroud, M. J. et al. Nesprin 1α2 is essential for mouse postnatal viability and nuclear positioning in skeletal muscle. J. Cell Biol. 216, 1915–1924 (2017)).

46. Levy, Y. et al. Prelamin A causes aberrant myonuclear arrangement and results in muscle fiber weakness. JCI Insight 3, (2018).

47. Metzger, T. et al. MAP and kinesin-dependent nuclear positioning is required for skeletal muscle function. Nature 484, 120–124 (2012).

48. Cristea, A. et al. Effects of aging and gender on the spatial organization of nuclei in single human skeletal muscle cells. Aging Cell 9, 685–697 (2010).

49. Meinke, P. et al. Muscular Dystrophy-Associated SUN1 and SUN2 Variants Disrupt Nuclear-Cytoskeletal Connections and Myonuclear Organization. PLoS Genet. 10, (2014).

50. Dubowitz, V., Sewry, C. & Oldfors, A. Muscle Biopsy: A Practical Approach. (2013).

51. Mcclelland, V. et al. Vici Syndrome Associated With Sensorineural Hearing Loss and Evidence of Neuromuscular Involvement on Muscle Biopsy. (2010). doi:10.1002/ajmg.a.33296

52. Taranum, S. et al. LINC complex alterations in DMD and EDMD/CMT fibroblasts. Eur. J. Cell Biol. 91, 614–628 (2012)).

53. Sanoudou, D. et al. Expression profiling reveals altered satellite cell numbers and glycolytic enzyme transcription in nemaline myopathy muscle. Proc. Natl. Acad. Sci. U. S. A. 100, 4666– 71 (2003).

54. Kotter, S., Andresen, C. & Kruger, M. Titin: Central player of hypertrophic signaling and sarcomeric protein quality control. Biol. Chem. 395, 1341–1352 (2014)).

55. Hu, L. Y. R., Ackermann, M. A. & Kontrogianni-Konstantopoulos, A. The sarcomeric M-region: A molecular command center for diverse cellular processes. Biomed Res. Int. 2015, (2015).

56. Jani, K. & Schöck, F. Molecular mechanisms of mechanosensing in muscle development. Dev. Dyn. 238, 1526–34 (2009)).

57. Belanto, J. J. et al. Microtubule binding distinguishes dystrophin from utrophin. Proc. Natl. Acad. Sci. U. S. A. 111, 5723–8 (2014)).

58. Sébastien, M. et al. Deletion of the microtubule-associated protein 6 (MAP6) results in skeletal muscle dysfunction. Skelet. Muscle 8, 1–14 (2018).

59. Khairallah, R. J. et al. Microtubules Underlie Dysfunction in Duchenne Muscular Dystrophy. Sci. Signal. 5, ra56-ra56 (2012).

60. O’Rourke, A. R. et al. Impaired muscle relaxation and mitochondrial fission associated with genetic ablation of cytoplasmic actin isoforms. FEBS J. 285, 481–500 (2018)).

61. Chapman, M. A. et al. Disruption of both nesprin 1 and desmin results in nuclear anchorage defects and fibrosis in skeletal muscle. Hum. Mol. Genet. 23, 5879–5892 (2014)).

62. Delhaas, T., Van der Meer, S. F. T., Schaart, G., Degens, H. & Drost, M. R. Steep Increase in Myonuclear Domain Size During Infancy. Anat. Rec. 296, 192–197 (2013)).

63. Ochala, J., Ravenscroft, G., Laing, N. G. & Nowak, K. J. Nemaline Myopathy-Related Skeletal Muscle ??-Actin (ACTA1) Mutation, Asp286Gly, Prevents Proper Strong Myosin Binding and Triggers Muscle Weakness. PLoS One 7, 1–6 (2012).

64. Ross, J. et al. Defects in glycosylation impair satellite stem cell function and niche composition in the muscles of the dystrophic largemyd mouse. Stem Cells 30, 2330–2341 (2012).

65. Ross, J. A. et al. SIRT1 regulates nuclear number and domain size in skeletal muscle fibers. J. Cell. Physiol. 7157–7163 (2018). doi:10.1002/jcp.26542

66. Ross, J. A. et al. Exploring the Role of PGC-1α in Defining Nuclear Organisation in Skeletal Muscle Fibres. 1270–1274 (2016). doi:10.1002/jcp.25678

