## Supplemental figures 1-3; supplemental tables 1-3 for "Impairments in contractility and cytoskeletal organisation cause nuclear defects in nemaline myopathy"

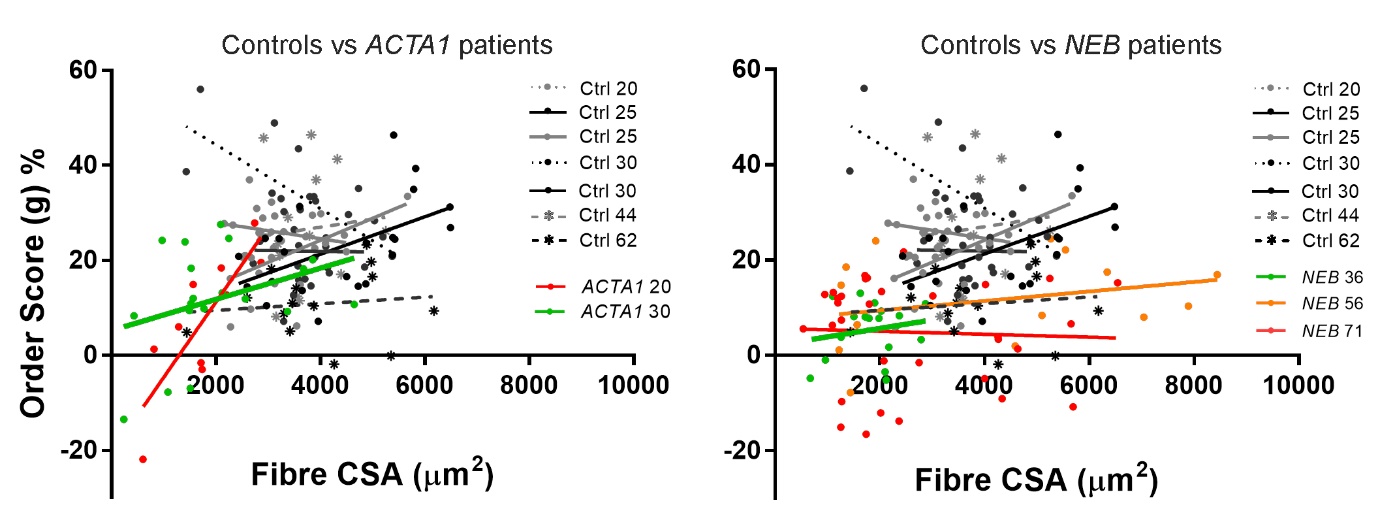

**Fig S1. Relationship between nuclear spacing and muscle fibre size.** Healthy control subjects and patients are denoted with their mutation and age. Order score (g), an algorithm to assess the regularity of nuclear spacing^27^; a lower score indicates more irregular spacing and more nuclear clustering. This parameter was plotted against fibre cross-sectional area (CSA). Individual data points represent an individual skeletal muscle fibre. Left, regression lines for control subjects versus *ACTA1* patients; right, regression lines for control subjects versus *NEB* patients. For most controls and patients, either no correlation, or a weak positive correlation between fibre CSA and order score was observed. One patient (*ACTA1 20*) showed a positive correlation (R^2^ = 0.68), indicating that larger fibres tended to be more ordered than small, although one control (*Ctrl 30*) showed a negative correlation (R^2^ = 0.47), suggesting the reverse relationship.

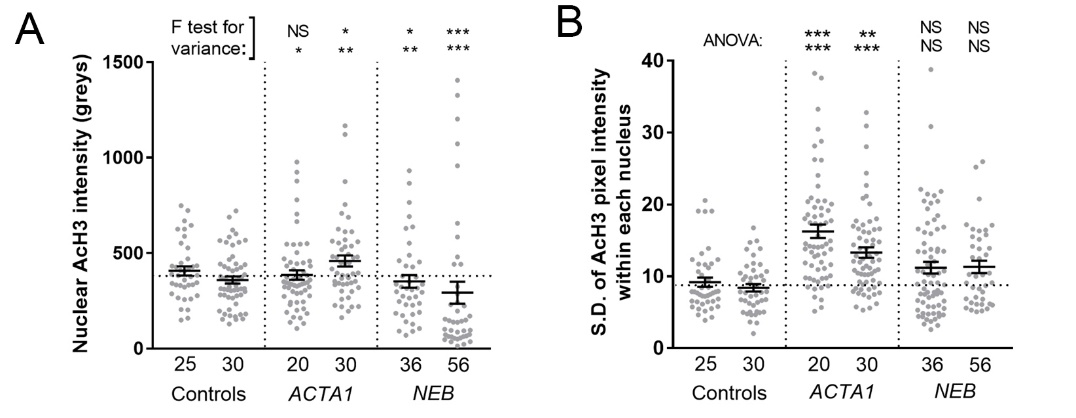

**Fig S2. Altered chromatin organisation in patients with nemaline myopathy (Related to Fig 3H-J).** **(A)** Mean acetylhistone H3 pixel intensity per nucleus (one data point per nucleus measured); F test for variance indicates that the variation in staining intensity between nuclei is significantly greater in patients than controls. **(B)** Standard deviation of pixel intensity within each nucleus, as a measure of staining variability within the nucleus (one data point per nucleus measured); patients frequently have more variable staining within each nucleus, possibly indicating irregularly packed regions of chromatin. 50+ nuclei were observed per subject across ~9 fibres, mean +/- SEM. * (P<0.05), ** (P<0.01), *** (P<0.001).

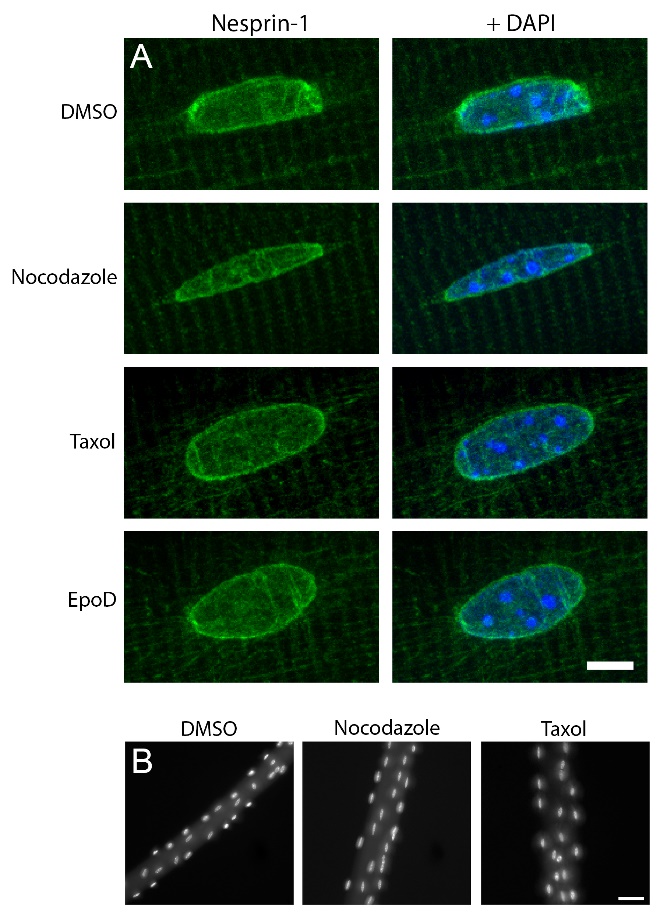

**Fig S3. Nesprin-1 localisation and myonuclear spacing is unaffected by microtubule perturbations.** **(A)** Typical myonuclei from mouse skeletal muscle fibres treated overnight with vehicle (DMSO), nocodazole, taxol or epothilone D. Nesprin-1 staining (green) and DAPI (blue). The localisation of nesprin-1 was not markedly affected by treatment with microtubule perturbing drugs. ~50 nuclei were observed per condition across 2-3 separate experiments. **(B)** Representative DAPI-stained images of muscle fibres treated with DMSO, nocodazole or taxol. No overt alterations to myonuclear spacing were observed in response to the drugs, even after 72 hours (~20 fibres observed across 2-3 experiments). Scale bars: 5μm (A); 50μm (B).

| **Gene** | **Age (yr)** | **M/F** | **Mutation (DNA)** | **Mutation (protein)** | **Source** |
| --- | --- | --- | --- | --- | --- |
| *ACTA1* | 20 | M | c.16G>A | E6K | Copenhagen, Denmark |
| *ACTA1* | 30 | F | c.841T>C | Y281H | Genoa, Italy |
| *NEB* | 36 | F | c.2836-2A>G and c.5763+5G>A | Mutation in splice site | Copenhagen, Denmark |
| *NEB* | 56 | M | c.17234C>T and c.2271_22713del | R5745X and K7571del | Copenhagen, Denmark |
| *NEB* | 71 | F | c.508-7T>A and c.19097G>T | Mutation in splice site; and S6366I | Helsinki, Finland |

**Table S1.** Patient muscle biopsy samples used for light microscopy.

| **Gene** | **Age (yr)** | **M/F** | **Mutation (DNA)** | **Mutation (protein)** | **Source** |
| --- | --- | --- | --- | --- | --- |
| *NEB* | 23 | M | c.11164C>T and c.19097G>T | R3722* (nonsense) and S6366I | Helsinki, Finland |
| *NEB* | 17 | M | c.22249A>C and c.8392-8395 duplication | T7417P and R2799L frameshift | Helsinki, Finland |
| *NEB* | 30 | F | c.508-7T>A and c.19097G>T | Mutation in splice site and S6366I | Helsinki, Finland |
| *NEB* | 2 | M | c.17737-2A>T and c.21315delA | Mutation in splice site and R7105 frameshift | Milan, Italy |
| *ACTA1* | 10 | M | c.841T>C | Y281H | Milan, Italy |
| *ACTA1* | 3 | M | c.235A>G | T79A | Milan, Italy |
| *ACTA1* | 11 weeks | F | c.796T>C | F266L | London, UK |

**Table S2.** Patient muscle biopsy samples used for electron microscopy.

| **Patient (mutation/age)** | **Chromatin density** | **Invaginations** | **Discontinuous and/or separation of nuclear membranes** | **Vacuolation of nuclear envelope** |
| --- | --- | --- | --- | --- |
| *NEB* 23 | ↑ | **+** | **++** | **+** |
| *NEB* 17 | ↓↓ | **++** | N | N |
| *NEB* 30 | ↑ | **+** | N | **+** |
| *NEB* 2 | ↑ | **+** | N | N |
| *ACTA1* 10 | ↓↓ | N | N | N |
| *ACTA1* 3 | ↓↓ | N | N | N |
| *ACTA1* 11 weeks | Only one nucleus image recorded | | | **+** |

**Table S3.** Ultrastructural observations in myonuclei of nemaline myopathy patients. Categorisation is based on criteria described in **Figure 3**. For chromatin density, ↑ denotes an increase and ↓ a decrease in density (respectively, an increase and reduction of heterochromatin). + indicates the presence of a given feature/observation. N denotes none observed. In all cases, two symbols indicates a particularly high incidence whereby the majority of observed myonuclei displayed the characteristic in question. 20 – 30 myonuclei observed in all patients, across multiple fibres and fields of view.
